# Reduced *ech-6* Expression Attenuates Fat-induced Premature Aging in *C. elegans*

**DOI:** 10.1101/2020.11.02.364760

**Authors:** Yasmine J. Liu, Arwen W. Gao, Reuben L. Smith, Georges E. Janssens, Daan M. Panneman, Aldo Jongejan, Michel van Weeghel, Frédéric M. Vaz, Melissa J. Silvestrini, Louis R. Lapierre, Alyson W. MacInnes, Riekelt H. Houtkooper

## Abstract

Deregulated energy homeostasis represents a hallmark of aging and results from complex gene-by-environment interactions. Here, we discovered that reducing the expression of the gene *ech-6* encoding enoyl-CoA hydratase remitted fat diet-induced deleterious effects on lifespan in *Caenorhabditis elegans*, while a basal expression of *ech-6* was important for survival under normal dietary conditions. Lipidomics revealed that supplementation of fat in *ech-6*-silenced worms had marginal effects on lipid profiles, suggesting an alternative fat utilization for energy production. Transcriptomics further suggest a causal relation between the lysosomal pathway, energy production, and the longevity effect conferred by the interaction between *ech-6* and high-fat diets. Indeed, enhancing energy production from endogenous fat by overexpressing lysosomal lipase *lipl-4* recapitulated the lifespan effects of high-fat diets on *ech-6*-silenced worms. Collectively, these results reveal that the gene *ech-6* modulates metabolic flexibility and may be a target for promoting metabolic health and longevity.

## INTRODUCTION

Lipids are an essential energy source required for numerous biological processes and tightly modulated to maintain organismal health (Smith et al., 2018; Wu et al., 2019). Dysfunctional lipid metabolism is associated with serious health problems including obesity, type 2 diabetes mellitus and cardiovascular diseases (Tilg et al., 2017; Wu et al., 2019; Yang et al., 2018). These pathological conditions have become epidemics of alarming proportions in many western countries where diets are enriched in fat (Gao et al., 2014; Mitchell et al., 2011). In addition, a growing body of evidence underscores the association between abnormal lipid metabolism and the aging process, in which a deficiency of fat breakdown occurs during aging and leads to fat accumulation in aged organisms (Gao et al., 2017; Houtkooper et al., 2011). Preventing fat accumulation, for instance by intermittent fasting, ameliorates aging and postpones the advent of aging-related metabolic disorders (Mattson et al., 2017).

Metabolic flexibility refers to a state of efficient switching between metabolic pathways (Smith et al., 2018). A flexible metabolism allows cells to adapt to fuel usage and to switch efficiently between nutrient sources depending on the environmental conditions (Smith et al., 2018). Gradual loss of this process during aging is a causative factor for increased susceptibility to aging-related metabolic disorders, yet the incidence and severity of these complex diseases vary considerably (Goodpaster and Sparks, 2017; Hunter, 2005). The underlying causes of the variability involve not only discrete genetic and environmental factors, but also the interactions between the two (Hunter, 2005; Williams and Auwerx, 2015). Gene-by-environment interactions (GxE) imply genetic predispositions that are differentially expressed depending on the environment and a genetic contribution to a specific phenotypic outcome that can be estimated when the environment is stable (Hunter, 2005; Williams and Auwerx, 2015). Previous forward and reverse genetic studies in model organisms have unraveled a number of susceptibility and resistance alleles that are relevant to complex traits and specific diseases (Andreux et al., 2012; Gao et al., 2018a; Kingsmore et al., 2007). However, our knowledge about the GxE pairs that explain the variability in the degree of metabolic flexibility and the rate of aging is still limited.

*Caenorhabditis elegans* is one of the most widely used model organisms in the aging field (Gao et al., 2018b; Olsen et al., 2006). Its fully sequenced genome and capability to respond to various dietary interventions facilitate the identification of GxE interactions influencing complex traits and the mechanistic delineation of molecular pathways (Hamilton et al., 2005; Hansen et al., 2007; Lee et al., 2003; Liu et al., 2019). Particularly, many of the core metabolic pathways that modulate aging in mammals are conserved in *C. elegans* (Houtkooper et al., 2010; Kenyon, 2010). For example, emerging evidence shows that lipid metabolism influences the lifespan of model organisms including *C. elegans* and depletion of regulators in fat metabolism shortens the lifespan of *C. elegans* (Taubert et al., 2006; Van Gilst et al., 2005). Nevertheless, the interaction between fat metabolism and aging is more convoluted, as different classes of long-lived mutant worms exhibit ranging levels of lipid content depending on the longevity signaling pathways involved, such as the insulin/insulin-like growth factor (IGF-1) pathway, the mTOR pathway, and the AMPK pathway (Gao et al., 2017; Heestand et al., 2013; Kimura et al., 1997; Shi et al., 2013).

In this study we aim to elucidate how genes and high-fat diets converge at the level of metabolic flexibility to affect longevity. We report that a gene encoding enoyl-CoA hydratase, *ech-6*, modulates worm lifespan in response to a dietary excess of fat. We show that high-fat feeding in wild-type animals leads to premature aging as a result of metabolic inflexibility induced by energy overload. Knockdown of *ech-6* under normal dietary conditions shortens lifespan, possibly caused by cellular energy crisis through suppressed mitochondrial function and compromised amino acid catabolism. Interestingly, knockdown of *ech-6* in combination with high-fat feeding protects animals against both fat diet- and *ech-6* RNAi-induced detrimental effect on lifespan. The underlying mechanism that accounts for the lifespan effect lies in the upregulated energy production and lysosome-related processes. The findings of this study indicate that the gene *ech-6* represents an adjustable factor that can be used for fine-tuning of the degree of metabolic flexibility in response to excessive dietary fat intake to modulate lifespan.

## RESULTS

### Reduced Expression of *ech-6* Prevents Accelerated Aging Caused by a Dietary Excess of Fat

To study the impact of dietary interventions on worm lifespan, particularly a high-fat diet intervention, we cultured wild-type N2 worms on plates containing soluble fat polysorbate 80 (P-80). We selected this compound, since (1) it contains a long-chain fatty acid, oleic acid, the most abundant fatty acid in adult *C. elegans* (Gao et al., 2017), and (2) it allows for uniform solubilization in the culture plates (Figure 1A). We found that worms exposed to a high-fat P-80 diet exhibited a significantly reduced lifespan (Figure 1B; Table S1), in line with the lifespan effect of high-fat diets in mice (Minor et al., 2011). Next, we conducted an RNAi-based lifespan screen to search for metabolic genes that could alter the susceptibility to the detrimental effects of this high-fat diet. One of the candidates that emerged was *ech-6* (enoyl-CoA hydratase 6; orthologue of mammalian *ECHS1*). Knockdown of *ech-6* protected worms against the decrease in lifespan upon P-80 supplementation (Figure 1C; Table S1). In addition, depletion of *ech-6* shortened lifespan (Figure 1C; Table S1), suggesting the requirement of a basal expression of *ech-6* for a normal lifespan. Taken together, these results suggest that *ech-6* interacts with high-fat diets to influence the rate of aging.

**Figure 1.**
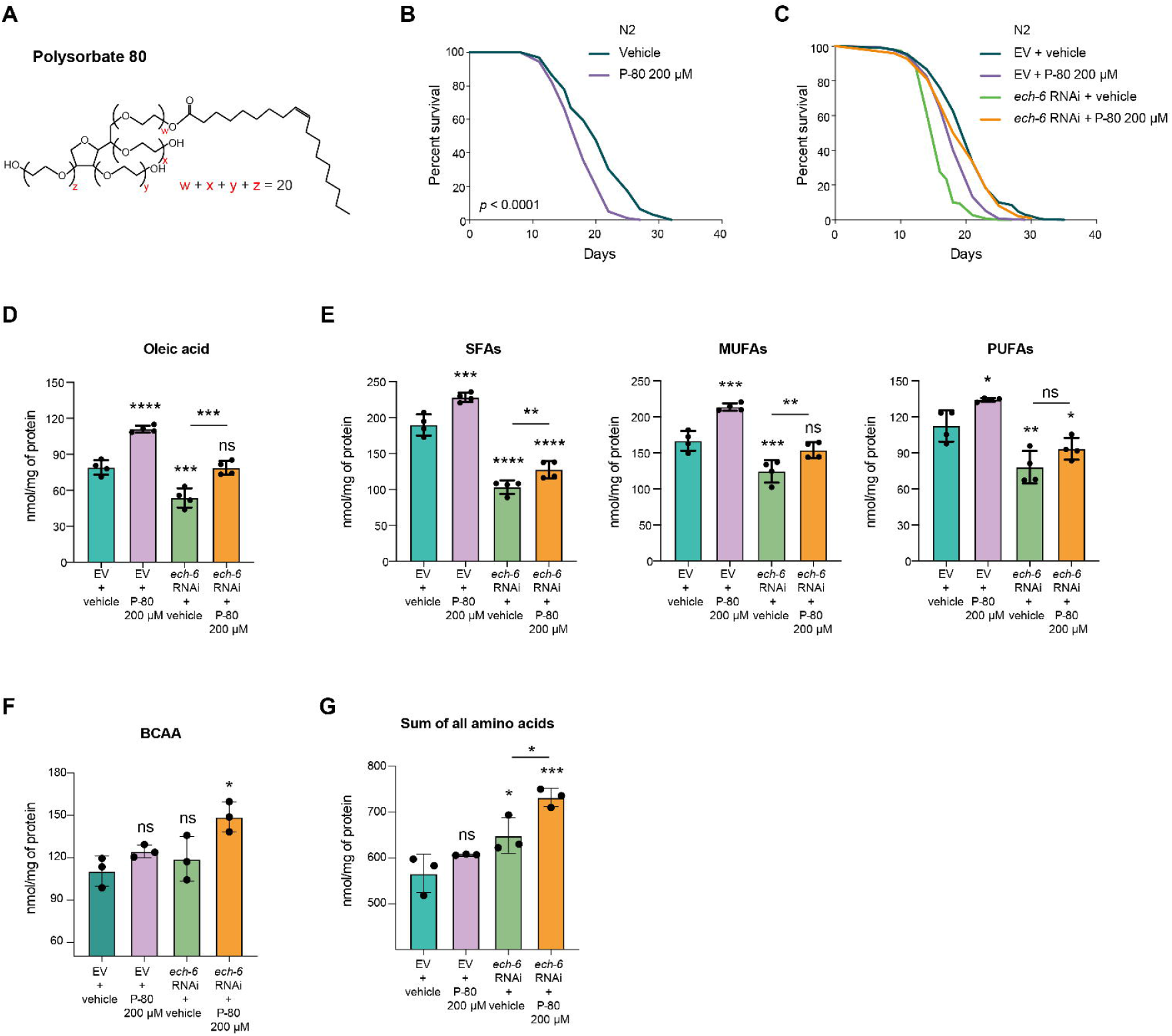
Knockdown of *ech-6* Antagonizes the Effects of a High-fat Diet on Lifespan, Oleic Acid Content and Fatty Acid Saturation Profile. (A) Chemical structure of polysorbate 80 (P-80). P-80 consists of a hydrophilic group of ethylene oxide polymers and oleic acid. (B) Survival curves of worms fed with 200 μM P-80 showing that worms live significantly shorter when subjected to a dietary excess of fat (*p* < 0.0001, log-rank test). (C) Survival curves of worms treated with *ech-6* RNAi or P-80 supplementation. Knockdown of *ech-6* protects worms against P-80 diet-induced lifespan decrease (*p* < 0.0001, log-rank test), while shortening lifespan under regular NGM dietary conditions (*p* < 0.0001, log-rank test). (D) Mass spectrometry (MS) quantification of oleic acid in worms exposed to the P-80 diet. P-80 supplementation increases the level of oleic acid in empty vector-treated wild-type (N2) worms while compensates for the reduced level caused by reduction of *ech-6* by RNAi. mean ± SD of 4 biological replicates. \*\*\**p* < 0.001; \*\*\*\**p* < 0.0001; ns, not significant; one-way ANOVA, Holm-Sidak correction. (E) MS quantification of saturated fatty acids (SFAs), monounsaturated fatty acids (MUFAs), and poly unsaturated fatty acids (PFAs). mean ± SD of 4 biological replicates. \**p* < 0.05; \*\**p* < 0.01; \*\*\**p* < 0.001; \*\*\*\**p* < 0.0001; ns, not significant; one-way ANOVA, Holm-Sidak correction. (F) UPLC-MS/MS quantification of branched-chain amino acids (BCAA). Knockdown of *ech-6* in combination with P-80 supplementation significantly increases the level of BCAA. mean ± SD of 3 biological replicates. \**p* < 0.05; ns, not significant; one-way ANOVA, Holm-Sidak correction. (G) Quantification of total amino acids. Knockdown of *ech-6* significantly increases the level of total amino acids. mean ± SD of 3 biological replicates. \**p* < 0.05; \*\*\**p* < 0.001; ns, not significant; one-way ANOVA, Holm-Sidak correction. An asterisk directly above a box refers to statistical significance compared to empty vector (EV)-treated wild-type (N2) worms fed a regular NGM diet, while an asterisk over a line indicates statistical significance compared to inhibition of *ech-6* by RNAi. See also Table S1 for lifespan data.

As oleic acid is a major component of P-80, we then asked whether feeding worms the high-fat diet would alter the endogenous level of oleic acid. To test this, we analyzed the fatty acid profile of worms exposed to the P-80 diet using a targeted metabolomics platform (Gao et al., 2017). Supplementation of P-80 to wild-type worms elevated the level of oleic acid (Figure 1D). Conversely, knockdown of *ech-6* decreased the level of oleic acid, which was increased to near-normal levels with the addition of P-80 (Figure 1D). As the saturation level of fatty acids was shown to be involved in longevity regulation in *C. elegans* (Han et al., 2017; O’Rourke et al., 2013), we next determined the effects of the P-80 diet on the degree of fatty acid saturation. Feeding wild-type worms with the P-80 diet significantly increased the levels of all three types of fatty acids including saturated fatty acids (SFAs), mono-unsaturated fatty acids (MUFAs) and poly-unsaturated fatty acids (PUFAs) (Figure 1E). Conversely, knockdown of *ech-6* significantly decreased the levels of SFAs, MUFAs, and PUFAs (Figure 1E). In addition, we found that supplementing *ech-6*-silenced worms with P-80 specifically increased both SFAs and MUFAs, while having no effect on PUFAs (Figure 1E). Taken together, these data show that depletion of *ech-6* and supplementation of P-80 exhibit reciprocal effects on the level of oleic acid and the saturation state of fatty acids.

As the gene *ech-6* has been annotated as an enoyl-CoA hydratase involved in the branched-chain amino acids (BCAA) breakdown which plays an important role in energy production in cells (Ferdinandusse et al., 2015; Watson et al., 2016), we determined the effects of reduced *ech-6* expression on amino acid profiles using ultra performance liquid chromatography tandem-mass spectrometry (UPLC-MS/MS) (Gao et al., 2017). Knockdown of *ech-6* did not change the level of BCAA, but significantly elevated the overall levels of amino acids of which increased alanine made up a major portion (Figures 1F, 1G and S1). Interestingly, we observed an increased level of BCAA when supplementing *ech-6*-silenced worms with P-80, whereas this did not occur in wild-type worms upon P-80 supplementation. These results imply that knockdown of *ech-6* impairs amino acids catabolism which is exacerbated by fat supplementation.

### High-fat Diets Affect the Lifespan upon *ech-6* Deficiency in a Dose- and Oleate-dependent Fashion

Next, we determined the dose effect of P-80 on lifespan by supplementing worms with P-80 at increasing concentrations including 100 μM, 200 μM and 400 μM. Wild-type worms had a comparable decrease in lifespan when fed with three P-80 concentrations (Figure 2A; Table S1). In contrast, subjecting *ech-6*-deficient worms to a P-80 diet at the respective concentrations yielded distinct lifespan outcomes in which 200 μM P-80 resulted in the greatest lifespan normalization (Figure 2B; Table S1).

**Figure 2.**
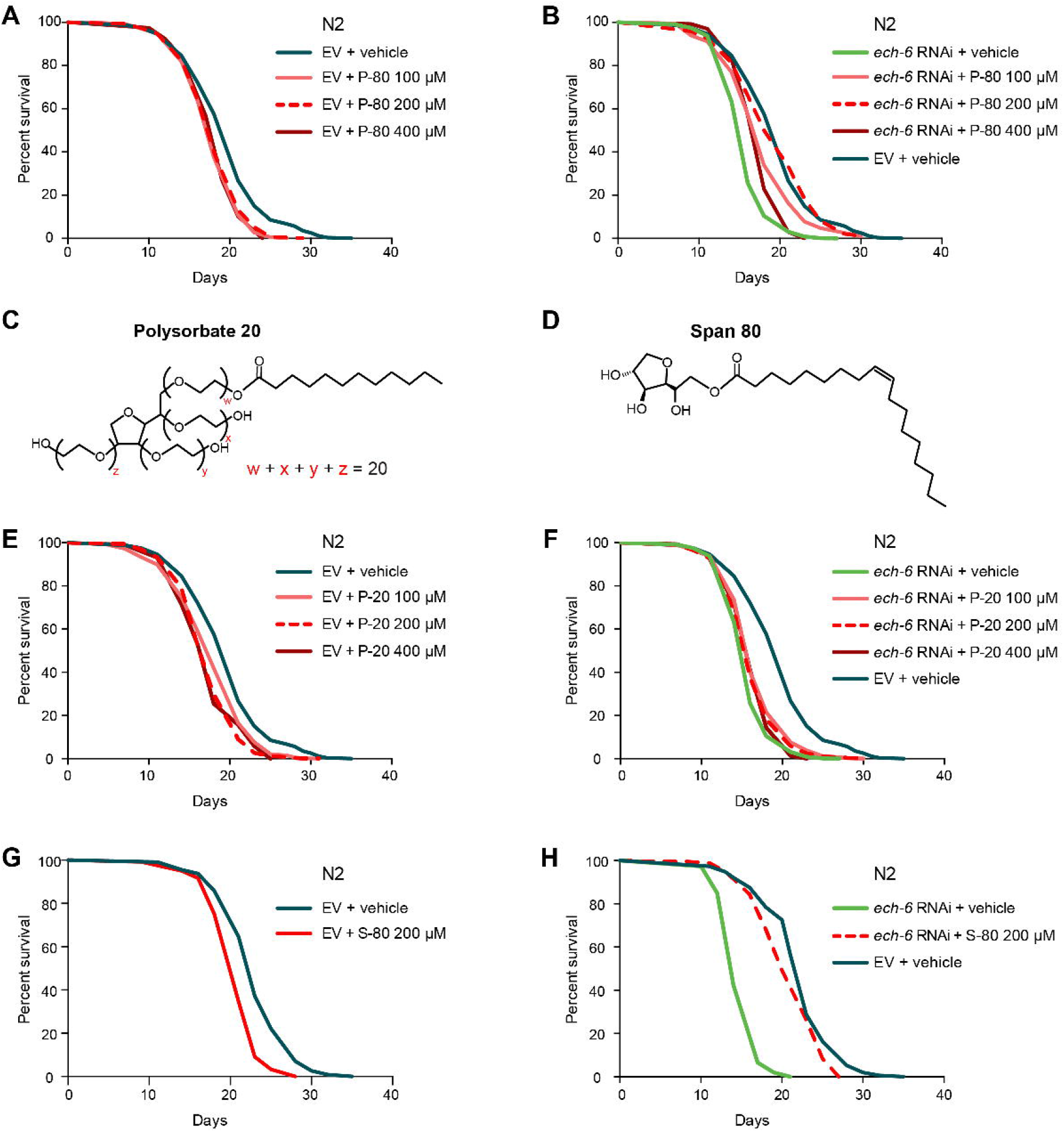
High-fat Diets Affect the Lifespan upon *ech-6* Deficiency in a Dose- and Oleic Acid-dependent Fashion. (A) Lifespan analysis showing that P-80 supplementation at doses ranging from 100 μM to 400 μM reduces lifespan (*p* < 0.0001, log-rank test). (B) Lifespan analysis of worms upon *ech-6* deficiency showing that feeding *ech-6*-deficient worms the P-80 diet at 200 μM results in a maximum restoration of lifespan (*p* = 0.564, comparing *ech-6* RNAi + P-80 200 μM to empty vector (EV) + vehicle, log-rank test). (C) Chemical structure of Polysorbate 20 (P-20). P-20 consists of a hydrophilic group of ethylene oxide polymers and a lauric acid. (D) Chemical structure of Span 80 (S-80). S-80 consists of a sorbitan monoester and an oleic acid. (E) Lifespan analysis of worms fed with P-20 showing that worms exposed to the P-20 diet at doses ranging from 100 μM to 400 μM lived significantly shorter (*p* < 0.0001, log-rank test). (F) Lifespan analysis of worms upon *ech-6* deficiency showing that P-20 supplementation at a concentration from 100 μM to 400 μM does not prevent *ech-6* RNAi from shortening lifespan (*p* < 0.0001, log-rank test). (G) Lifespan analysis of worms fed a S-80 diet showing that S-80 supplementation at 200 μM shortens lifespan (*p* < 0.0001, log-rank test). (H) Lifespan analysis of worms upon *ech-6* deficiency showing that S-80 supplementation at 200 μM protects worms against *ech-6* RNAi-induced lifespan decrease (*p* < 0.0001, log-rank test). See also Table S1 for lifespan data.

As the compound P-80 comprises a hydrophilic group of ethylene oxide polymers and an oleic acid moiety (Figure 1A), we asked which constituents of P-80 could be responsible for the lifespan phenotypes observed above. To test this, we made use of two other types of soluble fat, i.e., polysorbate 20 (P-20) and span 80 (S-80). P-20 consists of the same polar complex as P-80 but is bound to a saturated fatty acid with a shorter chain length, namely C12:0, lauric acid (Figure 2C). S-80 is another oleic acid derivative containing a smaller polar moiety (Figure 2D). Like the effects of P-80 on wild-type worms, supplementation of P-20 at 100 μM, 200 μM, and 400 μM shortened the lifespan of wild-type worms by a similar extent (Figure 2E; Table S1). In contrast to the effects of P-80 in worms with reduced *ech-6* expression, however, supplementation of P-20 at any of the three concentrations did not normalize their lifespan (Figure 2F; Table S1). Accordingly, we reasoned that rather than the hydrophilic polar complex of P-80, it is the oleic acid moiety that contributed to the restored lifespan of *ech-6*-deficient worms when fed with the high-fat P-80 diet. In line with this, supplementing *ech-6*-deficient worms with 200 μM S-80 extended their lifespan to near wild-type levels, while wild-type worms lived significantly shorter when fed with S-80 (Figures 2G and 2H; Table S1). Taken together, our results suggest that although different kinds of fat diets cause premature aging, *ech-6* specifically interacts with oleic acids to impinge on longevity.

### Knockdown of *ech-6* Suppresses Metabolic Activity and Energy-intensive Processes Including Growth and Mobility

Mitochondria provide for the majority of cellular energy via oxidative phosphorylation which consumes oxygen and generates ATP (Saraste, 1999). To further characterize whether knockdown of *ech-6* or addition of fat affected mitochondrial capacity for energy production, we measured mitochondrial respiration by monitoring oxygen consumption rate (OCR) using Seahorse respirometry (Koopman et al., 2016). Supplementation of P-80 to wild-type worms reduced the level of basal and maximal mitochondrial respiration (Figures 3A and 3B). In comparison, knockdown of *ech-6* also reduced mitochondrial respiration but to a greater extent than the P-80 diet, while feeding *ech-6*-deficient worms with P-80 did not show an additive effect (Figures 3A and 3B).

**Figure 3.**
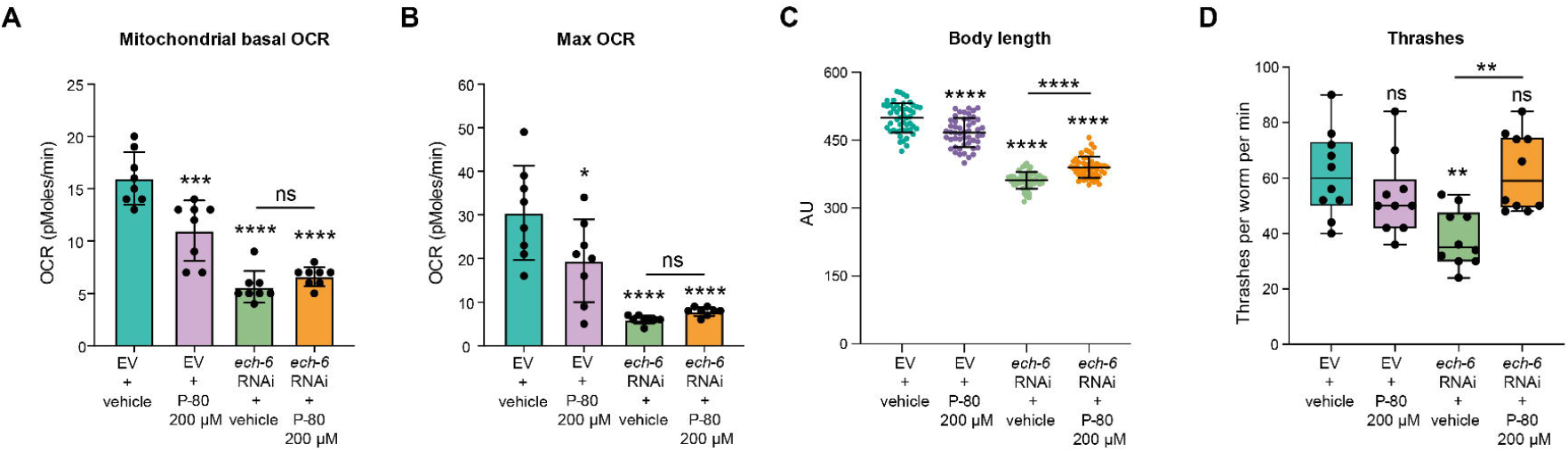
RNAi of *ech-6* Suppresses Metabolic Activity, Mobility, and Growth. (A-B) Mitochondrial basal and maximum oxygen consumption rate (OCR) in adult N2 worms at day 5 of adulthood. P-80 supplementation and *ech-6* RNAi decrease mitochondrial respiration rate at both the basal (A) and the maximal levels (B). Mean ± SD of n = 8 biological replicates. \*\*\**p* < 0.001; \*\*\*\**p* < 0.0001; one-way ANOVA, Holm-Sidak correction. (C) Body length of N2 worms at day 5 of adulthood. Mean ± SD of n = 50 animals. \*\*\*\**p* < 0.0001; one-way ANOVA, Holm-Sidak correction. P-80 supplementation ameliorates *ech-6* RNAi-mediated growth inhibition. (D) Thrashing of N2 worms at day 2 of adulthood exposed to *ech-6* RNAi and 200 μM P-80 diet, individually or in combination. P-80 supplementation to *ech-6*-deficient worms restores movement ability, measured by the frequency of thrashes per worm per minute. Mean ± SD of n = 10 animals. The bar graphs depict mean ± SD. \*\**p* < 0.01; ns, not significant; one-way ANOVA; Holm-Sidak correction. An asterisk directly above a box refers to statistical significance compared to empty vector (EV)-treated wild-type fed a regular NGM diet.

Growth and movement are two physiological processes that heavily rely on cellular energy supply where mitochondria play a paramount role (Saraste, 1999). We therefore asked whether body length and motility were compromised as a result of *ech-6* deficiency or high-fat feeding. Indeed, we found that wild-type worms subjected to the high-fat P-80 diet experienced a decrease in body length and remained small in size throughout life (Figure 3C), however their mobility was unaffected (Figure 3D). In comparison, knockdown of *ech-6* markedly reduced both the body size and the movement capacity (Figures 3C and 3D), while supplementing P-80 to *ech-6*-deficient worms significantly improved both parameters relative to that of *ech-6*-deficient worms (Figures 3C and 3D). Taken together, our observations show that depending on the expression level of *ech-6*, namely normal levels seen in wild type versus reduced levels by *ech-6* knockdown, the high-fat P-80 diet exhibits reciprocal effects on physiological functions involving growth and mobility.

### High-fat Diet-induced Alterations in Lipid Profiles are Diminished in the Context of *ech-6* Knockdown

Feeding wild-type worms with the high-fat P-80 diet shortened lifespan, decreased growth capacity, and reduced mobility, whereas P-80 supplementation upon knockdown of *ech-6* exhibited inverse effects (Figures 1C, 3C and 3D). We hence asked whether the way in which dietary fat was metabolized in the two genetic backgrounds may be relevant to the differences observed in the life history traits. To do this, we profiled lipids for wild-type and *ech-6-*deficient worms exposed to the high-fat P-80 diet using UPLC-MS-based lipidomics (Figure 4A). Across all worm samples, we detected 1151 distinct lipids belonging to 28 lipid species which can be further grouped into five main lipid classes consisting of diradylglycerol lipids, triradylglycerol lipids, glycerophospholipids, sterol lipids, and sphingolipids (Figures 4B and 4C; Table S2). Partial least squares discriminant analysis (PLS-DA) demonstrated that wild-type worms fed the P-80 diet were well separated from the control group fed a regular NGM diet by the first two principle components (Figure 4D). However, upon knockdown of *ech-6*, worms fed the P-80 diet overlapped with those fed a regular NGM diet, suggesting that the high-fat P-80 diet-induced lipid changes were diminished upon *ech-6* deficiency (Figure 4D).

**Figure 4.**
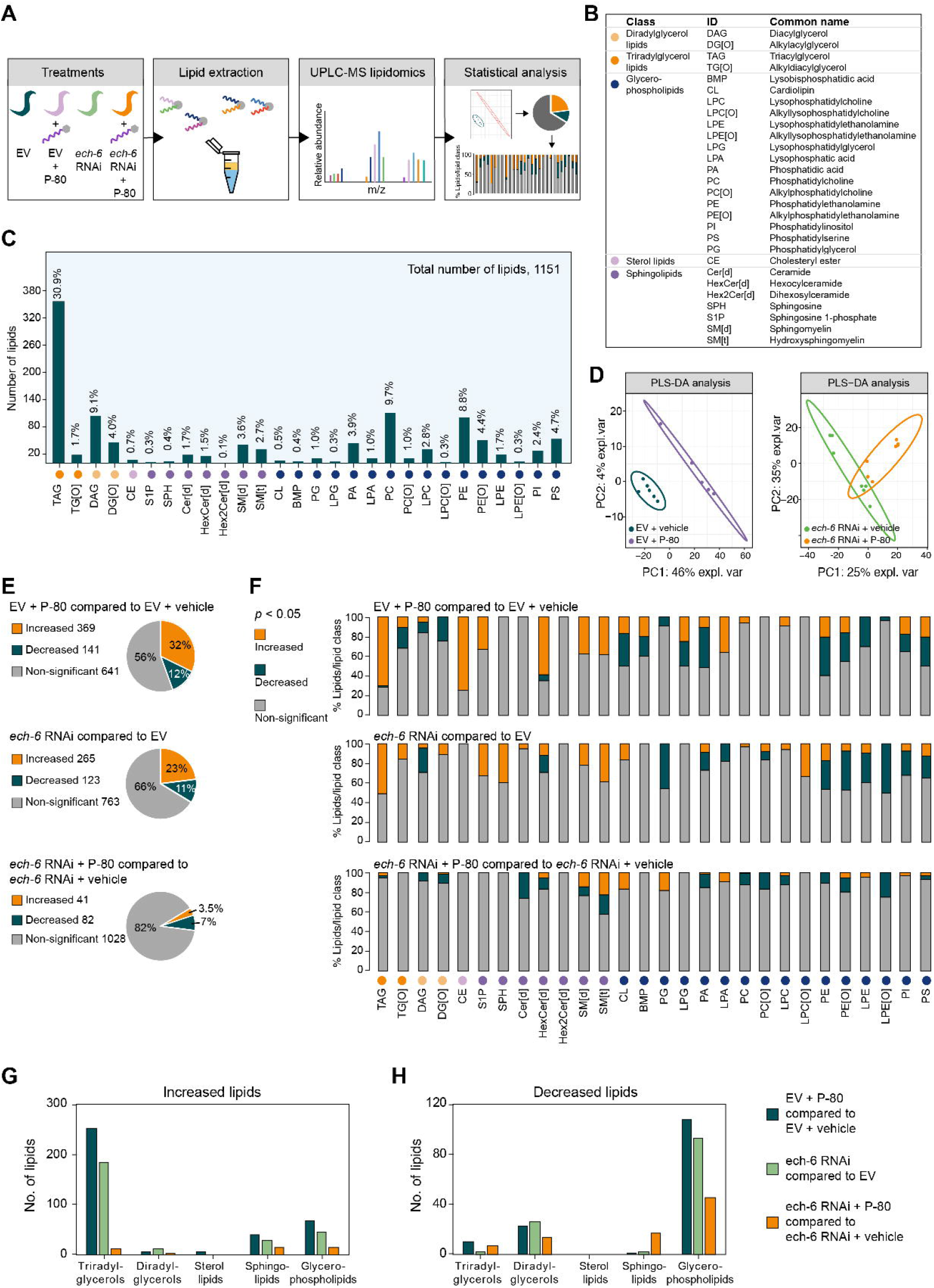
The Effects of High-fat Diets on Lipid Profiles are Abrogated upon *ech-6* Deficiency. (A) Experimental design. N2 worms were treated with *ech-6* RNAi or with P-80 from larval stage 1 and collected at day 1 adulthood for high-performance liquid chromatography-mass spectrometry (HPLC-MS)-based lipidomics. (B) Overview of worm lipids and distribution in the main lipid clusters. (C) The lipid composition of day-1 adult N2 worms. 28 lipid species and 1151 individual lipids were detected in worms. Triacylglycerides (TAG) make up the largest percentage (30.9%) of total lipids. (D) Partial least squares-discriminant analysis (PLS-DA) showing group separations based on significantly changed lipids by P-80 supplementation in empty vector (EV)- and *ech-6* RNAi-treated worms. (E) Pie charts depicting the percentages of increased and decreased lipids by P-80 supplementation or *ech-6* RNAi. A *p*-value < 0.05 was applied for determining statistical significance. (F) Percentage distribution of significantly changed lipids by P-80 supplementation across the lipid species in empty vector (EV)- and *ech-6* RNAi-treated worms. A *p*-value < 0.05 was applied to determine statistical significance. (G-H) The number of increased (G) and decreased (H) lipids in each lipid class upon P-80 supplementation or *ech-6* RNAi. The effects of P-80 supplementation on lipid profiles are significantly abolished upon knockdown of *ech-6* by RNAi. χ^2^ tests, *p* = 0, *p* = 0.2416, *p* = 1, *p* = 0, and *p* = 0.0421 when the number of lipids changed by P-80 supplementation including triradylglycerols, diradylglycerols, sterol lipids, sphingolipids, and glycerophospholipids in empty vector (EV)-treated worms was respectively compared to the corresponding lipid number in the context of *ech-6* deficiency. See also Figure S2 and Table S2.

We further confirmed this finding by analyzing the percentage of lipids showing significantly altered levels after supplementation of P-80. When wild-type worms were fed with P-80, 369 lipids were increased while 141 were decreased (Figure 4E; Table S2). Although knockdown of *ech-6* also elicited changes in a number of lipids relative to empty vector-treated wild-type controls (Figure 4E; Table S2), supplementation of P-80 to *ech-6*-deficient worms had a minor effect on lipid levels compared to *ech-6*-deficient worms fed a regular NGM diet, with only 41 increased and 82 decreased lipids (Figure 4E; Table S2). To further determine the patterns of the change for each lipid species, we analyzed the distribution of significantly altered lipids across all lipid species (Figure 4F; Table S2). Percentage distribution of both increased and decreased lipids within each lipid species showed that triacylglycerols (TAG), cholesteryl ester (CE), and sphingolipids including sphinganine 1-phosphate (S1P), hexocylceramide (HexCer[d]), sphingomyelin (SM[d]), and hydroxysphingomyelin (SM[t]) comprised almost solely increased lipids when wild-type worms were fed with P-80 (Figure 4F; Table S2). Under the same condition, the majority of decreased lipids were enriched in glycerophospholipids such as cardiolipin (CL), bis(monoacylglycero)phosphate (BMP), phosphatidylglycerol (PG), lysophosphatidylglycerol (LPG), phosphatidic acid (PA), phosphatidylethanolamine (PE), alkylphosphatidylethanolamine (PE[O]), lysophosphatidylethanolamine (LPE), phosphatidylinositol (PI), and phosphatidylserine (PS) (Figure 4F; Table S2). In addition, the lipid classes triradylglycerols and glycerophospholipids possessed the greatest number of lipids affected by P-80 supplementation in wild-type worms, among which 251 triradylglycerols were increased while 108 glycerophospholipids were decreased (Figures 4G and 4H; Table S2). These results imply that supplementation of P-80 to wild-type worms does not merely give rise to accumulation of storage lipids (triradylglycerols), but also profoundly change the profile of membrane lipids such as the glycerophospholipids. Knockdown of *ech-6* altered lipid profiles in a similar fashion as did P-80 supplementation, i.e., triradylglycerols and sphingolipids comprised primarily increased lipids, while the majority of decreased lipids were enriched in glycerophospholipids (Figures 4F-4H; Table S2). However, in contrast to the widespread effects of P-80 on wild-type worms, supplementation of P-80 to *ech-6-*deficient worms had greatly reduced effects on all lipid classes compared to the lipid levels in *ech-6*-deficient worms (Figures 4F-4H; Table S2). These results suggest that excessive fat intake in *ech-6*-deficient worms may be used as fuel for energy production, for instance through fatty acid β-oxidation, rather than being converted into triacylglycerols for storage or into glycerophospholipids for membrane synthesis.

To determine whether the effects of a fat diet on lipid profiles were specific to oleic acid-enriched fat, we measured lipid changes occurring upon the addition of lauric acid-enriched fat P-20. Feeding wild-type worms with P-20 changed the pattern of overall lipid profiles in a way analogous to the effects of P-80, including a large number of increased triradylglycerols and decreased glycerophospholipids (Figures S2A-S2D; Table S2). In comparison, supplementation of P-20 to *ech-6*-deficient worms affected a reduced number of triradylglycerols (Figures S2B and S2C; Table S2), however, other lipid classes including sphingolipids and glycerophospholipids were changed in *ech-6*-deficient worms to a similar extent as seen in wild-type worms (Figures S2B-S2D; Table S2). Taken together, these results show that knockdown of *ech-6* renders animals resistant to overall lipid alterations caused by oleic acid-rich fat diets like P-80.

### Energy Production and Lysosomal Functions Constitute the Core Set of Upregulated Biological Processes upon High-fat Feeding

To understand the underlying mechanisms accounting for the longevity effect resulting from the interaction between *ech-6* and high-fat diets, we conducted RNA sequencing (RNA-seq) analyses. Wild-type worms subjected to the P-80 diet had large numbers of transcripts differentially expressed (Figure 5A; Table S3), among which 1816 and 1256 transcripts were down- and upregulated (Figure 5B; Table S3). Upon knockdown of *ech-6*, the total number of transcripts affected by P-80 supplementation were significantly decreased (Figure 5A), with only 831 and 558 remaining down- and upregulated (Figure 5B). Next, we asked to what extent the transcriptional changes induced by P-80 supplementation in wild-type worms were different from those upon knockdown of *ech-6*. To address this, we compared up- and downregulated transcripts by P-80 supplementation in wild-type worms to those in *ech-6*-silenced worms (Figure 5C), which can be categorized into three groups: (G1) up- or downregulated transcripts exclusively present in *ech-6*-silenced worms; (G2) up- or downregulated transcripts co-existing in both wild-type and *ech-6*-silenced worms; (G3) up- or downregulated transcripts specific to wild-type worms (Figure 5C). Function enrichment analysis of the transcripts from each group revealed that catabolic pathways such as carbohydrate catabolic process, aminoglycan catabolic process and polysaccharide catabolic process were exclusively overrepresented in the upregulated genes in *ech-6*-silenced worms upon P-80 supplementation (G1) (Figure 5D; Table S3). In contrast, anabolic pathways including lipid biosynthesis and carboxylic acid biosynthesis were enriched among the upregulated genes exclusively present in wild-type worms (G3) (Figure 5D; Table S3). On the other hand, biological processes involved in energy homeostasis and lipid transport were enriched among the commonly upregulated transcripts in both wild-type and *ech-6*-silenced worms (G2) (Figure 5D; Table S3). Taken together, these results suggest that genes involved in energy homeostasis and lipid transport represent a core set of high-fat responsive genes, regardless of the expression of *ech-6*, while lipid biosynthesis and carbohydrate catabolism are conditionally upregulated, depending on *ech-6*.

**Figure 5.**
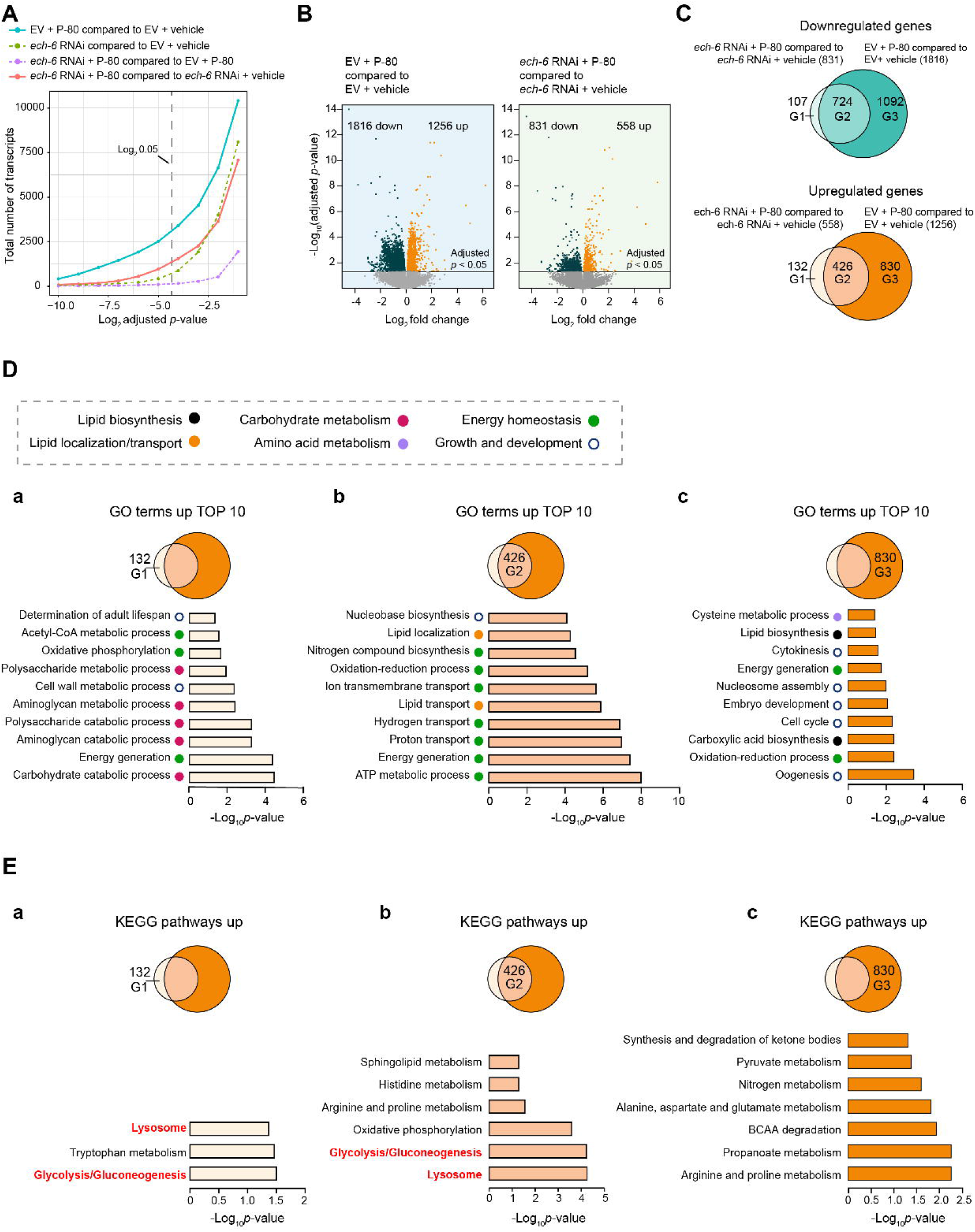
High-fat Feeding Provokes Activation of a Core Set of Biological Processes Related to Energy Production and Lysosomal Functions. (A) The total number of transcripts change with adjusted *p*-values upon high-fat feeding or depletion of *ech-6* by RNAi. An adjusted *p*-value was calculated for each transcript in the respective comparisons between P-80-fed wild-type worms versus those fed a regular NGM diet, between *ech-6*-deficient versus wild-type worms, between P-80-fed *ech-6*-deficient versus P-80-fed wild-type worms, and between P-80 fed *ech-6*-deficient worms versus those fed a regular NGM diet. P-80 supplementation to empty vector (EV)-treated wild-type worms induces a great number of differentially expressed transcripts and the transcriptional effects of P-80 supplementation are reduced in the context of *ech-6* RNAi, as shown by the comparison between *ech-6* RNAi + P-80 and *ech-6* RNAi + vehicle. Adjusted *p*-values (Benjamini-Hochberg method) < 0.05 was used to determine the statistical significance of the difference. (B) Volcano plots depicting differentially expressed transcripts induced by P-80 supplementation in empty vector (EV)- and *ech-6* RNAi-treated worms. Adjusted *p*-values (Benjamini-Hochberg method) < 0.05 was applied to define significantly up- and downregulated genes. (C) Venn diagrams comparing overlaps between genes down- and upregulated by P-80 supplementation in empty vector (EV)- and *ech-6* RNAi-treated worms. G1, differentially down- or upregulated genes exclusively in *ech-6*-deficient worms. G2, commonly down- or upregulated genes in empty vector (EV)- and *ech-6* RNAi-treated worms. G3, down- or upregulated genes exclusively in empty vector (EV)-treated worms. (D) Gene ontology enrichment analysis of upregulated genes belonging to G1, G2, and G3, respectively. GO terms were considered significantly enriched for a modified Fisher’s Exact *p*-value < 0.05 (an EASE score). (E) KEGG pathway enrichment analysis of upregulated genes belonging to G1, G2, and G3, respectively. Pathways were considered significantly enriched for a modified Fisher’s Exact *p*-value < 0.05 (an EASE score). See also Figure S3 and Table S3.

To further determine the specific pathways in which each group of upregulated genes was involved, we performed KEGG pathway enrichment analysis. Pathways involving lysosome, glycolysis/gluconeogenesis, and oxidative phosphorylation were identified as the top 3 most significantly enriched pathways among the commonly upregulated transcripts present in both wild-type and *ech-6*-deficient worms (G2) (Figure 5E; Table S3). Moreover, lysosome and glycolysis/gluconeogenesis were again overrepresented for the upregulated transcripts exclusively present in *ech-6*-deficient worms (G1) (Figure 5E; Table S3), highlighting the importance of these pathways upon *ech-6* deficiency in combination with high-fat feeding. In addition to the two pathways, only tryptophan metabolism was found as another upregulated pathway in the context of *ech-6* RNAi (Figure 5E; Table S3). As opposed to the specific upregulation of lysosome and glycolysis/gluconeogenesis upon knockdown of *ech-6* (Figure 5E; Table S3), a number of pathways regulating various aspects of metabolism were enriched among the upregulated transcripts exclusively in wild-type worms upon P-80 supplementation (G3) (Figure 5E; Table S3). Among those were pathways involving amino acid metabolism such as BCAA degradation, and alanine, aspartate and glutamate metabolism, short chain fatty acid propanoate metabolism, and pyruvate and ketone body metabolism (Figure 5E; Table S3). Functional enrichment analysis of the downregulated transcripts revealed few enrichments, with the exception of processes related to growth and development, which were found in the downregulated gene clusters commonly present in both wild-type and *ech-6*-deficient worms (G2) (Figure S3A) and in those exclusively in wild-type worms (G3) (Figure S3B; Table S3). Taken together, these results suggest that supplementation of P-80 rather upregulates energy production and lysosome-related processes upon *ech-6* deficiency, as opposed to a broader metabolic influence in wild-type worms involving stimulation of other processes such as lipid biosynthesis, various amino acid metabolic pathways, and pyruvate and ketone body metabolism.

### Upregulation of *lipl-4* Underlies High-fat Diet-mediated Lifespan Effects upon *ech-6* Deficiency

Given that the lysosomal pathway emerged as one of the most significantly upregulated pathways by P-80 supplementation, in particular upon *ech-6* deficiency (Figure 5E), we asked whether upregulation of lysosomal genes could be relevant to the effects of the P-80 diet on the lifespan of *ech-6*-deficient worms. To address this, we profiled the differential expression of lysosomal genes and found that the majority was significantly upregulated after P-80 supplementation to *ech-6*-deficient worms (Figure 6A). To search for candidate genes that account for P-80-mediated longevity effect in *ech-6*-deficient worms, we focused on the top 10 upregulated lysosomal genes (Figure 6B). We found that 4 out of 10 genes encode lysosomal lipases including *lipl-1*, *lipl-3*, *lipl-4*, and *lipl-7*, while the other 6 genes are involved in sphingolipid metabolism and proteolysis, including *asah-1*, *asm-2*, and *asm-3* for sphingolipid metabolism, and *cpr-1, cpr-4,* and *cpr-5* for proteolysis (Figure 6B). Lipases break down complex fat molecules such as triglycerides into their component fatty acids and glycerol molecules through hydrolysis of ester bonds so as to provide substrates for energy production or storage (Zechner et al., 2017). Given the fact that the ester bond linking oleate to the hydrophilic group in the P-80 compound is similar to that in triglycerides, we hypothesized that the upregulated lipases in *ech-6*-deficient worms after P-80 supplementation may facilitate the digestion of P-80 for an ultimate breakdown. The lysosomal lipase LIPL-4 is the most well-characterized in relation to fat metabolism and longevity in *C. elegans* and overexpression of *lipl-4* has been shown to promote fat mobilization and adjust mitochondrial activity (Folick et al., 2015; Lapierre et al., 2011; Ramachandran et al., 2019; Wang et al., 2008). Therefore, we asked whether overexpression of *lipl-4* could mimic the effect of P-80 supplementation on the lifespan of *ech-6*-deficient worms by promoting endogenous fat mobilization. To do this, we developed an integrated version of a transgenic strain that expresses *lipl-4* under its own promoter (Lapierre et al., 2011; Wang et al., 2008) and characterized their lipid profiles using UPLC-MS-based lipidomics. We found that overexpression of *lipl-4* in *ech-6*-deficient worms significantly reduced the level of almost all triglyceride species and that of total triglycerides content (Figures 6C and 6D; Table S2), indicating that these fats have been released by overexpressing *lipl-4* for energy production. Given that feeding *ech-6*-deficient worms with P-80 restored the lifespan to a near-normal level, we next examined whether enhancing endogenous fat utilization through overexpression of *lipl-4* could similarly benefit the lifespan of *ech-6*-deficient worms. Indeed, overexpressing *lipl-4* fully rescued the lifespan of *ech-6*-deficiency (Figure 6E; Table S1). Altogether, our data support that enhanced lysosomal lipolysis restores lipid mobilization and lifespan, similar to P-80 supplementation, when ech-6 function is impaired.

**Figure 6.**
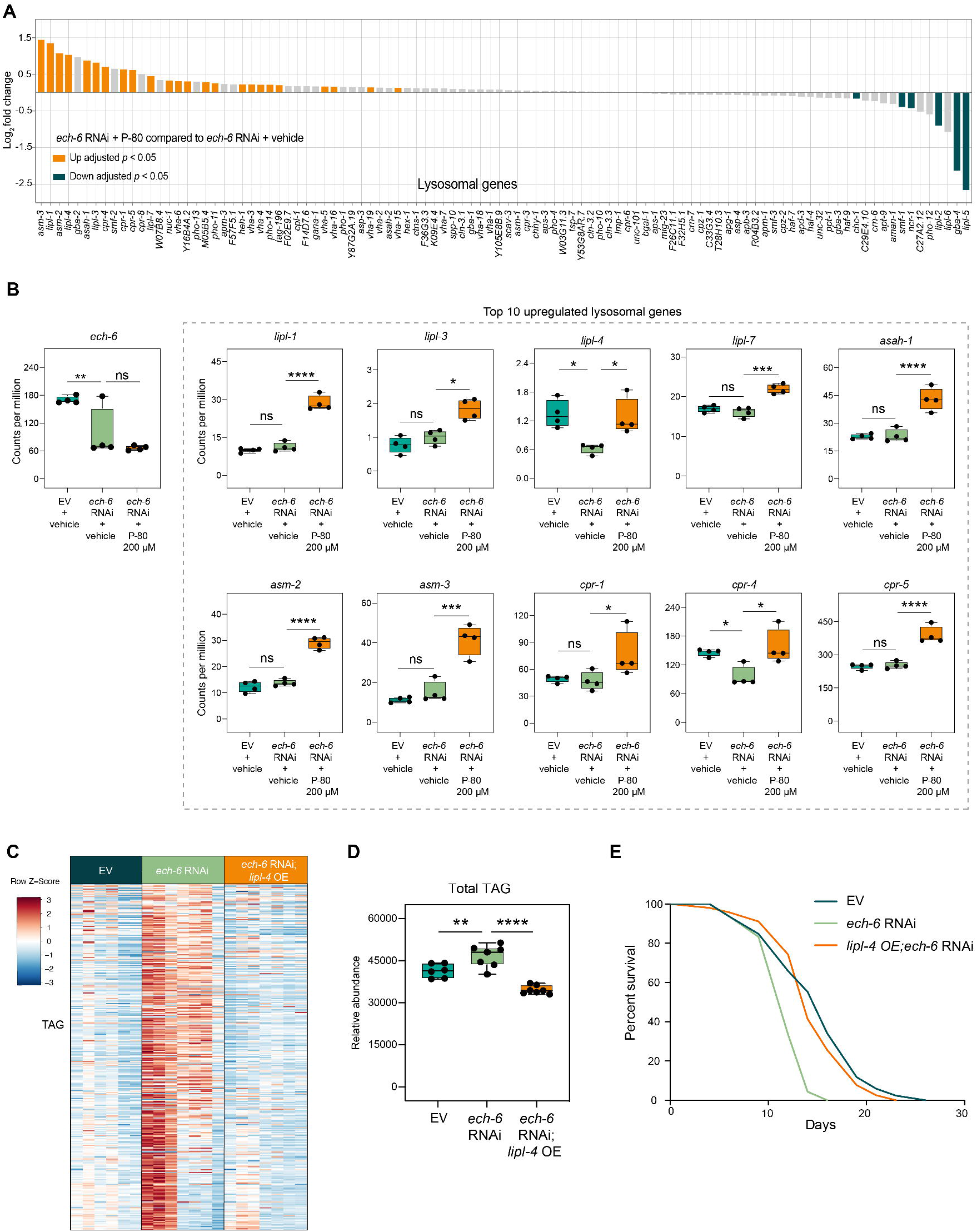
Overexpression of *lipl-4* Promotes Fat Mobilization and Restores the Lifespan of *ech-6*-deficient Worms. (A) Expressional changes of lysosome genes induced by P-80 supplementation in *ech-6*-deficient worms as compared to *ech-6*-deficient worms fed a regular NGM diet. 24 and 6 lysosome genes were respectively up- and downregulated. Adjusted *p*-values (Benjamini-Hochberg method) < 0.05 was considered significantly different. (B) Selection of the top 10 significantly upregulated transcripts in (A). 4 out of 10 upregulated transcripts encode lysosome lipases including *lipl-1*, *lipl-4*, *lipl-3*, and *lipl-7*. \**p* < 0.05; \*\**p* < 0.01; \*\*\**p* < 0.001; \*\*\*\**p* < 0.0001; ns, not significant; one-way ANOVA; Adjusted *p*-values (Benjamini-Hochberg method) < 0.05 was considered significantly different. (C) Heatmap showing that overexpression of *lipl-4* decreases the level of triacylglycerols (TAG) upon knockdown of *ech-6* as compared to that in worms treated with *ech-6* RNAi alone. The lipid species of triacylglycerols were ordered from top to bottom by acyl chain length. (D) Quantification of the total amount of TAG showing that overexpression of *lipl-4* decreases the total TAG content upon knockdown of *ech-6* as compared to that in worms treated with *ech-6* RNAi alone. \**p* < 0.05; \*\**p* < 0.01; one-way ANOVA; Holm-Sidak correction. (E) Survival curves of worms overexpressing *lipl-4* subjected to *ech-6* RNAi showing that *lipl-4* overexpression restores the lifespan of *ech-6*-depleted worms to near-wild type levels (*p* = 0.2631, comparing *lipl-4* OE;*ech-6* RNAi to empty vector (EV), log-rank test). See also Table S1 for lifespan data and Table S2 for triacylglycerol measurement.

## DISCUSSION

In this study, we uncover the interactions between *ech-6* and high-fat diets in regulation of lifespan in *C. elegans*. Upon high-fat feeding, decreasing the expression of *ech-6* rendered worms resistant to the deleterious effects of a high-fat diet on lifespan. However, under normal dietary conditions, knocking down *ech-6* shortened lifespan. The gene *ech-6* encodes a protein in BCAA catabolism. Knocking down *ech-6* resulted in amino acid accumulation and inhibited metabolic activity, thereby causing energy crisis and suppressing physiological activities that are dependent on energy homeostasis. Supplementation of fat to *ech-6*-deficient worms prevented the lifespan decrease and partially restored the normal function of the physiological processes. UPLC-MS-based lipidomics revealed that although high-fat feeding profoundly altered lipid profiles in wild-type worms particularly by increasing storage lipids triacylglycerols, the effect of high-fat diets on lipid content was abolished in *ech-6*-deficient worms. Using a transcriptomics approach followed by confirmation experiments, we identified that upregulation of lysosomal activity for instance by overexpression of lysosomal lipase *lipl-4* underlies the lifespan response to fat diets upon *ech-6* deficiency. Together, these results emphasize a novel role of ECH-6 in regulating metabolic flexibility by virtue of upregulation of lysosomal pathway in response to dietary fat overload and in turn affecting lifespan.

In this work, we unraveled a new connection between *ech-6* and high-fat diets in modulating aging through their interactive effects on energy metabolism in *C. elegans*. As illustrated in the model shown in Figure 7, we suggest that worms grown on a normal diet are metabolically flexible, which enables them to switch efficiently between available nutrient sources and to maintain energy homeostasis (Pang and Curran, 2014). This in turn leads to a normal lifespan (Figure 7A). When worms are exposed to excessive dietary fat, this metabolic balance is interrupted, leading to reduced mitochondrial function, increased lipid biosynthesis and accumulated lipids, and in turn causing premature aging and shortened lifespan (Figure 7A). In contrast, this situation is completely changed when the expression of *ech-6* is reduced in these worms. *ech-6* encodes an enoyl-CoA hydratase based on the homology with the human ECHS1 and is likely involved in the degradation of branched-chain amino acids valine and isoleucine (Ferdinandusse et al., 2015; Wanders et al., 2012). We hence propose that upon knockdown of *ech-6—*when energy from branched-chain amino acids becomes unavailable— these worms go through an adaptive state that increases reliance on other nutrient sources such as fat (Figure 7B). This hypothesis is supported by our findings that worms with reduced expression of *ech-6* do not accumulate fat when exposed to a high-fat diet (Figures 4E-4H). This adaptation allows them to cope better with the load of dietary fat and thereby to prevent premature aging induced by a high-fat diet (Figure 7B). Knockdown of *ech-6* on the other hand leads to a metabolic crisis driven by disrupted BCAA degradation and suppressed mitochondrial respiration, in turn resulting in accelerated aging. However, this metabolic crisis is ameliorated by either supplying additional oleate-enriched fat to *ech-6*-deficient worms or by boosting endogenous fat mobilization for energy production through overexpression of *lipl-4* (Figure 7B).

**Figure 7.**
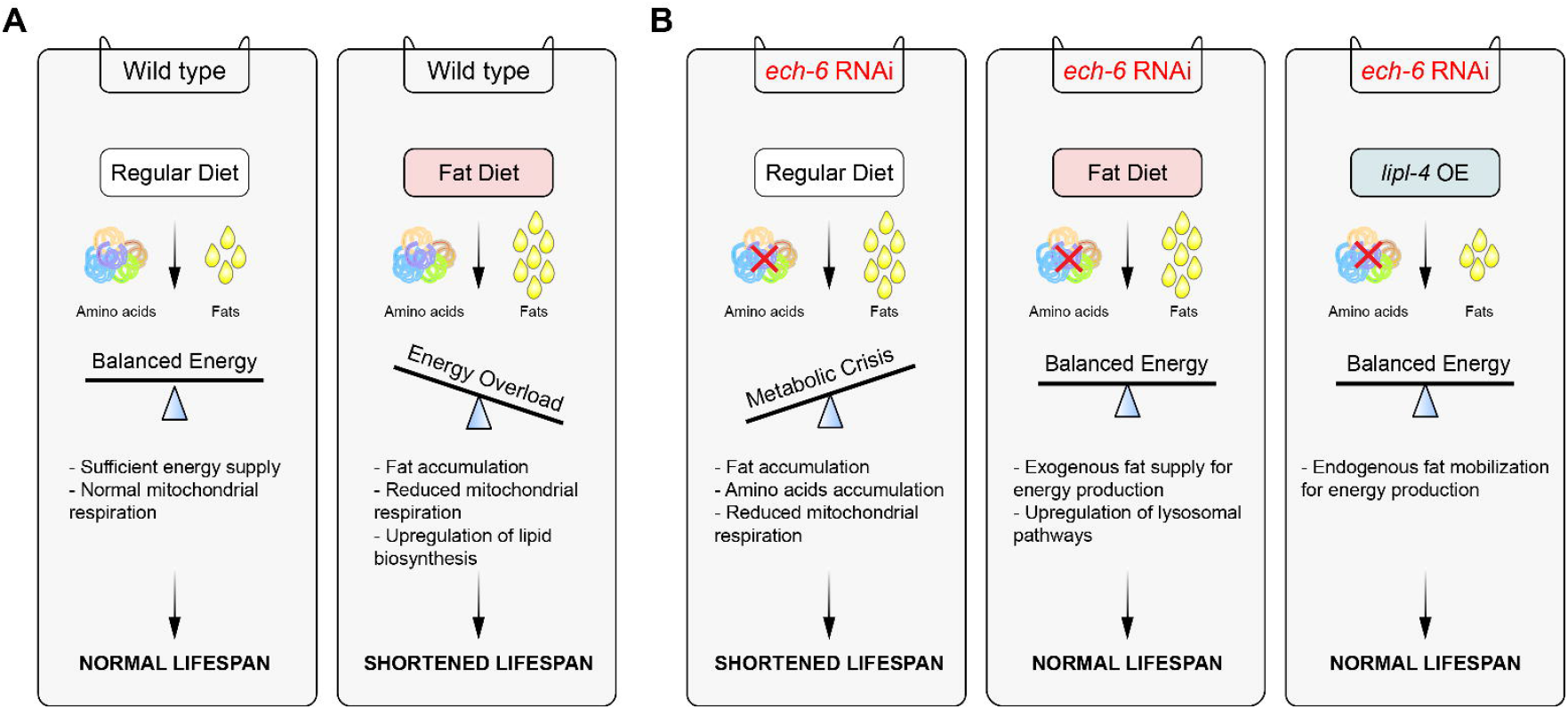
Model for Hypothetic Interactions between *ech-6, lipl-4,* a High-fat Diet, and Lifespan. (A) On a regular diet, worms have sufficient energy supply from mitochondria to maintain a homeostatic balance between energy production and energy consumption. As such, worms display a normal lifespan. However, when worms are exposed to excessive dietary diet, lipid overload disturbs energy homeostasis, suppresses mitochondrial respiration, causes fat accumulation, and ultimately leads to a shortened lifespan. (B) Knockdown of *ech-6* engenders metabolic crisis under normal dietary conditions by disrupting branched-chain amino acid degradation, inhibiting mitochondrial respiration, and resulting in fat and amino acids accumulation, which in turn causes insufficient energy production and lifespan reduction. This disrupted energy balance can be restored by either supplying dietary fat to *ech-6*-depleted worms or by enhancing endogenous fat mobilization for energy production through overexpression of *lipl-4*.

We found that knockdown of *ech-6* shortened lifespan which was prevented by supplementation of fat consisting of oleic acid rather than of lauric acid. We speculate that the different lifespan effects between the two high-fat diets could be due to two reasons: (1) P-20 and P-80 differ in their constituent fatty acids, oleic acid versus lauric acid and the latter of which releases around 1.5 times more ATP when fully broken down. Feeding *ech-6*-deficient worms with either high-fat diet had marginal effects on the level of TAG compared to the effects observed in wild-type worms upon high-fat feeding. This suggests that the additional fat in *ech-6*-deficient worms was primarily catabolized through fatty acid β-oxidation. Therefore, due the lower capacity of P-20 to produce energy compared to that of P-80, it is possible that the concentration we tested in our study may not suffice to extend the lifespan of *ech-6*-deficent worms; (2) supplementing P-80 to *ech-6*-deficient worms had greatly reduced effects on glycerophospholipid profiles, whereas this attenuation was not observed when *ech-6*-deficient worms were fed with the P-20 diet. Glycerophospholipids are major components of membranes, alterations of which will inevitably affect various aspects of cell functions and lifespan.

We found that knockdown of *ech-6* substantially reduced mitochondrial respiration. A closer investigation of this gene from a previous study showed the function of the encoded protein ECH-6 in degradation of propionate through the propionate shunt pathway (Watson et al., 2016), in addition to the initially predicated function in the breakdown of BCAA in *C. elegans* (Watson et al., 2013). Perturbations in the propionate shunt pathway, especially through depletion of *ech-6,* could result in accumulation of propionate and intermediates such as acrylyl-CoA—the substrate of ECH-6—to toxic levels (Watson et al., 2016). Therefore, we presume that knockdown of *ech-6* incurs damages to mitochondria partially due to the production of those toxic metabolites.

Taken together, our work identifies *ech-6* as a novel regulator of metabolic flexibility modulating the susceptibility towards dietary fat overload. Although a substantial loss of *ech-6* expression or function is deleterious to lifespan in both *C. elegans* and humans, it is appealing to speculate that moderate suppression of this gene such as through pharmacological approaches could improve metabolic flexibility towards fat diet without compromising health and survival. We believe that this project opens up new avenues to elucidate longevity genes involved in GxE interactions.

## ACKNOWLEDGMENTS

Work in the Houtkooper group is financially supported by an ERC Starting grant (no. 638290), and a VIDI grant from ZonMw (no. 91715305), and a grant from the Velux Stiftung (no. 1063). A.W.G. was supported by an AMC PhD Scholarship. A.W.M. is supported by E-Rare-2, the ERA-Net for Research on Rare Diseases (ZonMW #40-44000-98-1008). G.E.J. is supported by a Federation of European Biochemical Society (FEBS, https://www.febs.org) long-term fellowship and a VENI grant from ZonMw (no. 09150161810014, https://www.zonmw.nl). The authors thank the Caenorhabditis Genetics Center at the University of Minnesota for providing *C. elegans* strains. The authors also thank Lodewijk IJlst and Simone W. Denis for advice on the manuscript.

## AUTHOR CONTRIBUTIONS

Y.J.L., A.W.G., R.L.S., A.W.M., and R.H.H. conceived and designed the project. Y.J.L., A.W.G., R.L.S., R.K., D.M.P., and M.J.S. performed the experiments. Y.J.L., G.E.J. and A.J. performed RNAseq bioinformatics. M.v.W. and F.M.V. performed and interpreted metabolomics and lipidomics. L.R.L. provided worms strains and advice. Y.J.L., A.W.G., A.W.M. and R.H.H. wrote the manuscript with contributions from all other coauthors.

## STAR METHODS

- KEY RESOURCES TABLE
- LEAD CONTACT AND MATERIALS AVAILABILITY
- EXPERIMENTAL MODEL AND SUBJECT DETAILS

○ *C. elegans* Strains and Bacterial Feeding Strains
- METHOD DETAILS

○ Nematode Growing Conditions and RNAi Experiments
○ Supplementation of Polysorbate 80 (P-80), Span 80 (S-80) or Polysorbate 20 (P-20)
○ Lifespan Measurements
○ Analysis of Fatty Acid Profile Using Targeted Mass Spectrometry-based Platform
○ Amino Acid Extraction and UPLC-MS/MS Analysis
○ Analysis of Worm Body Length
○ Thrashing Assay
○ Oxygen Consumption Rate Analysis
○ One-phase Lipidomic Extraction and Lipidomics in *C. elegans*
○ Isolation of mRNA
○ Library Preparation
○ Read Mapping, Statistical Analyses, and Data Visualization
○ Functional Annotation of Gene Sets
- QUANTIFICATION AND STATISTICAL ANALYSIS
- DATA AND CODE AVAILABILTY

## LEAD CONTACT AND MATERIALS AVAILABILITY

This study did not generate new unique reagents. Further information and requests for resources and reagents should be directed to and will be fulfilled by the Lead Contact, Riekelt H. Houtkooper.

## EXPERIMENTAL MODEL AND SUBJECT DETAILS

### *C. elegans* Strains and Bacterial Feeding Strains

The *C. elegans* N2 (Bristol) and *E. coli* OP50 was obtained from the *Caenorhabiditis* Genetics Center (CGC). The LIPL-4 OE strain (LRL21) is an integrated version of the GR1971 strain (mgEx779[lipl-4p/K04A8.5p::lipl-4/K04A8.5::SL2::gfp + myo-2p::mcherry]) (Wang et al., 2008) that was backcrossed 4 times to wild-type. RNAi bacterial clones are *E. coli* HT115 strains, including the clone to knock down *ech-6 (*T05G5.6) which was derived from Ahringer library (Kamath et al., 2003). The *ech-6* RNAi clone was confirmed by sequencing. In this study, RNAi experiments initiated at larval stage 1.

### METHOD DETAILS

#### Nematode Growing Conditions and RNAi Experiments

*C. elegans* were cultured and maintained on nematode growth media (NGM) at 20°C. Worms of each strain were cultured on plates seeded with OP50 strain *Escherichia coli*. In RNAi experiments, synchronized N2 at L1 stage were obtained by alkaline hypochlorite treatment of gravid adults and transferred onto NGM plates (containing 2 mM IPTG and 25 mg/mL carbenicillin) seeded with *E. coli* HT115 containing empty vector or *ech-6* RNAi bacteria.

#### Supplementation of Polysorbate 80 (P-80), Span 80 (S-80) or Polysorbate 20 (P-20)

Pure P-80 (Merck KGaA), S-80 (Croda) or P-20 (Merck KGaA) was added into the liquid NGM agar medium before autoclaving to obtain a final concentration of 100 μM, 200 μM, or 400 μM.

#### Lifespan Measurements

Worm lifespan was performed as described previously (Liu et al., 2019). In short, a synchronized population of L1 worms of each strain was obtained as described above and seeded onto NGMi (containing 2 mM IPTG and 25 mg/mL carbenicillin) plates or NGMi plates containing different concentrations of P-80, S-80 or P-20. After worms reached the last larval stage L4, these worms were then transferred onto NGMi (or NGMi + P-80/S-80) plates containing 10 μM 5-FU. Worms were scored every other day. Worms that did not react to gentle stimulation were scored as dead. Worms that crawled off the plates or displayed a protruding vulva phenotype were censored.

### Analysis of Fatty Acid Profile Using Targeted Mass Spectrometry-based Platform

Fatty acid extraction was performed as mentioned in our previous study (Gao et al., 2017). A synchronized population of 2000 L1 worms were cultured on plates seeded with *E. coli* HT115 or *ech-6* RNAi bacteria for 2.5 days until reaching young adult stage. Worms were then collected by washing off the culture plates with M9 buffer followed by three times of washing with dH_2_O. Worm pellet was transferred to a 2 mL Eppendorf tube using a glass pipette followed by snap-freezing in liquid N_2_ and freeze-drying overnight. Worm lysate was generated using a TissueLyser II (Qiagen) (adding a 5 mm steel bead and ice-cold 0.9% NaCl solution to a dried worm pellet) for 2×2.5 min at a frequency of 30 times/sec. The lysate was homogenized further by a tip sonication (energy level: 40 joule; output:8watts) on ice water. BCA assay was used for protein quantification and used for sample normalization. A 150 μg worm protein lysate was transferred in a 4 mL fatty acid free glass vial followed by adding 1 mL of freshly prepared mixture of pure acetonitrile/37% HCl (ratio 4:1, v/v) to the lysate. Deuterium-labeled internal standard mixture (5.04 nmol d5-C18:0, 2.52 nmol d4-C24:0, and 0.25 nmol d4-C26:0) was added to each vial and then the fatty acid samples were hydrolyzed by incubating at 90°C for 2 h. After the samples were cooled down to room temperature, 2 mL hexane was added to each vial and sample were mixed by vortexing for 5 sec. After 1 min centrifugation at 1000 *g*, the upper layer was transferred to a fatty acid free glass tube and evaporated at 30°C under a flow of nitrogen. Fatty acid pellets were dissolved in 150 μL final solution (chloroform-methanol-water, 50:45:5, v/v/v) containing 0.0025% aqueous ammonia and then transferred to a Gilson vial for ESI-MS analysis.

### Amino Acid Extraction and UPLC-MS/MS Analysis

Amino acid extraction and UPLC-MS/MS were performed as previously described in our study (Gao et al., 2017). In brief, 1 mL of 80% acetonitrile plus 20 μL of internal standard mixture containing 68◻nmol d4-alanine, 44◻nmol d3-glutamate, 40◻nmol d3-leucine, 28◻nmol d5-phenylalanine, 34◻nmol d8-valine, 34◻nmol d3-methionine, 26◻nmol d4-tyrosine, 22◻nmol d5-tryptophan, 46◻nmol d3-serine, 48◻nmol d7-proline, 24◻nmol d7-arginine, 28◻nmol d5-glutamine, 32◻nmol d4-lysine, 26◻nmol ^13^C-citrulline, 28◻nmol d6-ornithine, 42◻nmol d10-isoleucine, and 46◻nmol d3-aspartate) were added to worm lysate and homogenized by vortexing in a 2 mL Eppendorf tube. Samples were centrifuged for 10 min at 4°C at 16,000 g and the supernatant was transferred to a 4 mL glass vial and evaporated under nitrogen stream at 40°C. Subsequently, the residue was dissolved in 220 μL of 0.01% (v/v in MQ water) heptafluorobutyric acid for UPLC-MS/MS analysis.

### Analysis of Worm Body Length

After exposure to the various conditions, worms were washed with M9, and imaged with a Leica M295 microscope. Measurements were performed with Image J freeware (W.S. Rasband, U.S.A. National Institutes of Health, Bethesda, Maryland, USA, http://rsb.info.nih.gov/ij/, 1997–2012). Measurements were performed on 50 worms for each condition.

### Thrashing Assay

After being treated with *ech-6* RNAi bacteria and 200 μM P-80, 10 adult worms at day 2 were used for the thrashing assay of each condition. In brief, a single worm was placed in a drop of M9 buffer on a clean glass slide and allowed to acclimatize for 30 seconds. The frequency of body bends was counted for 30 seconds as described previously (Nazir et al., 2010).

### Oxygen Consumption Rate Analysis

Worm oxygen consumption rate (OCR) was measured using the Seahorse XF96 (Seahorse Bioscience) as previously described (Koopman et al., 2016). In brief, worms were cultured on plates with different conditions and washed off with M9 buffer after reaching day 1 adulthood. After three times of additional washing with M9 buffer, worms were transferred in 96-well Seahorse plates and OCR was measured six times. FCCP and sodium azide treatments were performed at a final concentration of 10 μM and 40 mM, respectively. Both mitochondrial (basal) and maximum OCR was measured for each condition.

### One-phase Lipidomic Extraction and Lipidomics in *C. elegans*

Worms were synchronized at L1 and subjected to *ech-6* RNAi bacteria, 200 μM P-20, or 200 μM P-80 for 2.5 days until reaching day-1 adult stage. Lipidomics was performed as described (Gao et al., 2017; Molenaars et al., 2020). Briefly, samples containing approximately 2000 worms were lyophilized in 2 mL tubes. The samples were homogenized and lipids extracted in 1:1 (v/v) methanol:chloroform using water bath sonication for 10 min in presence of internal standards, these were: Bis(monoacylglycero)phosphate BMP(14:0)_2_ (0.2 nmol), Cardiolipin CL(14:0)_4_ (0.1 nmol), Cholesterol ester CE(16:0)-D7 (2.5 nmol), Diacylglyceride DG(14:0)_2_ (0.5 nmol), Lysophosphatidicacid LPA(14:0) (0.1 nmol), Lysophosphatidylcholine LPC(14:0) (0.5 nmol), Lysophosphatidylethanolamine LPE(14:0) (0.1 nmol), Lysophosphatidylglycerol LPG(14:0) (0.02 nmol), Phosphatidic acid PA(14:0)_2_ (0.5 nmol), Phosphatidylcholine PC(14:0)_2_ (2 nmol), Phosphatidylethanolamine PE(14:0)_2_ (0.5 nmol), Phosphatidylglycerol PG(14:0)_2_ (0.1 nmol), Phosphatidylinositol PI(8:0)_2_ (0.2 nmol), Phosphatidylserine PS(14:0)_2_ (5 nmol), Ceramide phosphocholines SM(d18:1/12:0) (2.125 nmol), Triacylglyceride TG(14:0)_3_ (0.5 nmol) (Avanti Polar Lipids, Alabaster, AL). After centrifugation at 16.000xg at 4°C for 10 mins, the supernatant was collected in glass vials and dried under a stream of nitrogen gas at 45°C and reconstituted in 150 μL 1:9 (v/v) methanol:chloroform. Chromatography was performed on a Dionex Ultimate 3000 binary UHPLC (Thermo Scientific) and on normal- and reversed phase polarity. MS data were acquired using negative and positive ionization using continuous scanning over the range of m/z 150 to m/z 2000. Data were analyzed using an in-house developed metabolomics pipeline written in the R programming language (http://ww.r-project.org). In brief, it consisted of the following five steps: (1) pre-processing using the R package XCMS, (2) identification of metabolites, (3) isotope correction, (4) normalization and scaling and (5) statistical analysis. All reported lipids were normalized to corresponding internal standards according to lipid class, as well as total protein content in samples, determined using a PierceTM BCA Protein Assay Kit. Lipid identification has been based on a combination of accurate mass, (relative) retention times, and the injection of relevant standards.

### Isolation of mRNA

For isolation of total mRNA, approximately 500 day-1 adult worms were collected in quadruplicates for each treatment. In brief, worm pellets were homogenized in TRIzol (Invitrogen) with a 5mm steel metal bead and the isolation was continued according to manufacturer’s protocol. For RNAseq, genomic DNA residues were removed using RNase-Free DNase (QIAGEN) and samples were cleaned up with the RNeasy MinElute Cleanup Kit (QIAGEN).

### Library Preparation

RNA libraries were prepared and sequenced with the illumina platform by Genome Scan (Leiden, Netherlands). Samples were processed for Illumina using the NEBNext Ultra Directional RNA Library Prep Kit (NEB #E7420) according to manufacturer’s description. Briefly, rRNA was depleted using the rRNA depletion kit (NEB# E6310). Subsequently, a cDNA synthesis was performed in order to ligate with the sequencing adapters. Quality and yields after sample preparation were measured with the Fragment Analyzer (Agilent). Sizes of the resulting products was consistent with the expected size distribution (a broad peak between 300–500 bp). Clustering and DNA sequencing using the Illumina cBot and HiSeq 4000 was performed according to manufacturer’s protocol with a concentration of 3.0 nM of DNA. HiSeq control software HCS v3.4.0, image analysis, base calling, and quality check was performed with the Illumina data analysis pipeline RTA v2.7.7 and Bcl2fastq v2.17.

### Read Mapping, Statistical Analyses, and Data Visualization

Reads were subjected to quality control FastQC (Andrews, 2010) trimmed using Trimmomatic v0.32 (Bolger et al., 2014) and aligned to the *C. elegans* genome obtained from Ensembl, wbcel235.v91 using HISAT2 v2.0.4 (Kim et al., 2015). Counts were obtained using HTSeq (v0.6.1, default parameters) (Anders et al., 2015) using the corresponding GTF taking into account the directions of the reads. Statistical analyses were performed using the edgeR (Robinson et al., 2010) and limma/voom (Ritchie et al., 2015) R packages. All genes with more than 2 counts in at least 4 of the samples were kept. Count data were transformed to log2-counts per million (logCPM), normalized by applying the trimmed mean of M-values method (Robinson et al., 2010) and precision weighted using voom (Law et al., 2014). Differential expression was assessed using an empirical Bayes moderated t test within limma’s linear model framework including the precision weights estimated by voom (Law et al., 2014; Ritchie et al., 2015). Resulting *p*-values were corrected for multiple testing using the Benjamini-Hochberg false discovery rate. Genes were re-annotated using biomaRt using the Ensembl genome databases (v91). Data processing was performed using R v3.4.3 and Bioconductor v3.5. The RNA-seq data are available at the Gene Expression Omnibus (GEO) Database under the ID GSE157701 and token: kvqlgiccbpczbct.

### Functional Annotation of Gene Sets

Gene sets were analyzed for functional enrichments using the DAVID bioinformatics resource version 6.7 (Huang da et al., 2009). Functional annotation clustering was performed using DAVID defined default settings incorporating gene sets from Gene Ontologies (biological process, cellular component, and molecular function), functional categories (including Clusters of Orthologous Groups (COG) ontologies, SP keywords, UP seq features), pathways (including KEGG), and protein domains (including INTERPRO PIR superfamily and SMART). The measured transcriptome was used as the background dataset for the enrichment tests and ReviGo (Supek et al., 2011) was used to eliminate redundant GO terms for purposes of visualization and summarization. Venn diagrams were generated using BioVenn (Hulsen et al., 2008).

### QUANTIFICATION AND STATISTICAL ANALYSIS

Statistical details, the number of biological replicates and any other forms of quantification present are specified in the respective figure legends and results section. Statistical analyses for lifespan, fatty acids, amino acids, mitochondrial respiration, body length and mobility were performed using the Prism 7 software (GraphPad Software, La Jolla, CA, USA). All comparisons of means were accomplished using a one-way ANOVA and two sample unpaired t-test as indicated in the corresponding figure legends. The significance *p*-values were adjusted to correct for multiple testing using the Holm-Sidak method. All other statistics were specified in each respective methods section unless otherwise noted. Gene expression was considered differential relative to respective control groups using adjusted *p*-values (Benjamini-Hochberg method) lower than 0.05. Lipid level was considered altered relative to respective control groups using *p-*values lower than 0.05. For lifespan studies, survival curves were calculated using the log-rank (Mantel-Cox) method.

### DATA AND CODE AVAILABILTY

The accession number for the *C. elegans* RNA sequencing data reported in this paper is GEO: GSE157701 and GEO reviewer token is kvqlgiccbpczbct. The processed and normalized RNAseq data is also available as Table S3.

