## Supplemental Figures for "Reduced *ech-6* Expression Attenuates Fat-induced Premature Aging in *C. elegans*"

**SUPPLEMENTAL FIGURES AND LEGENDS**

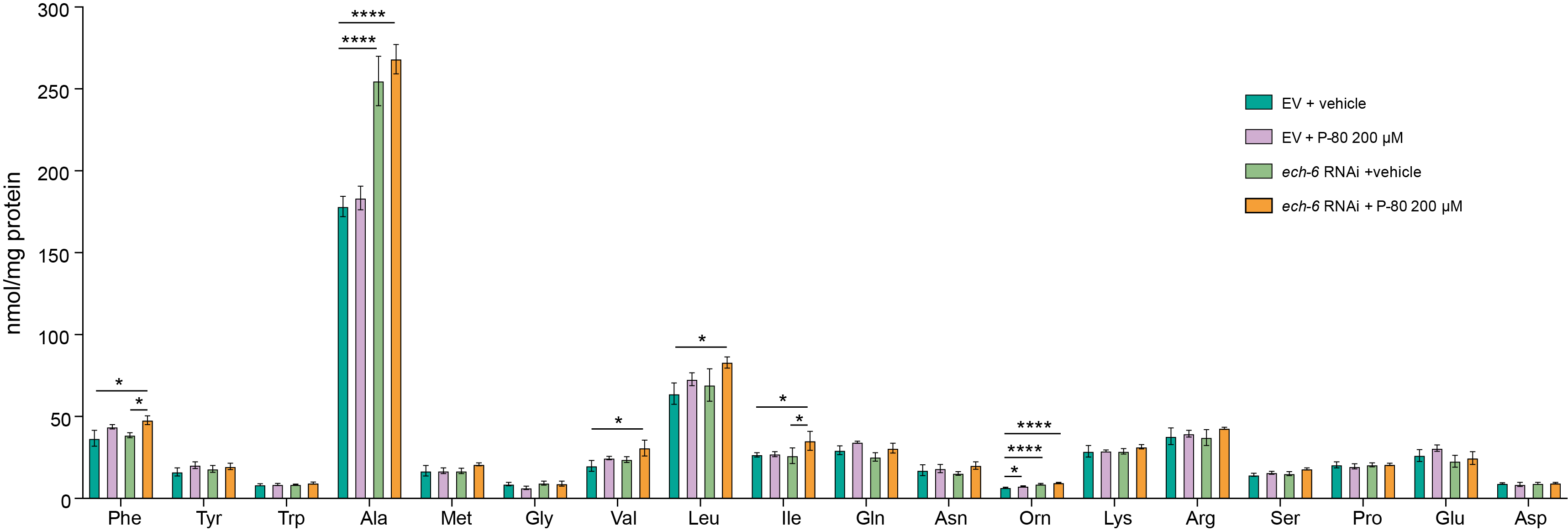

**Figure S1.** Amino Acid Profiles upon *ech-6* Depletion and High-fat Feeding, Individually or in Combination

Knockdown of *ech-6* significantly increases the levels of valine and ornithine, while supplementation of P-80 to *ech-6*-silenced worms increases the levels of branched-chain amino acids (valine, leucine, and isoleucine), phenylalanine, and ornithine as compared to the empty vector (EV) controls. **p* < 0.05; ***p* < 0.01; ****p* < 0.001; *****p* < 0.0001; one-way ANOVA; Holm-Sidak correction.

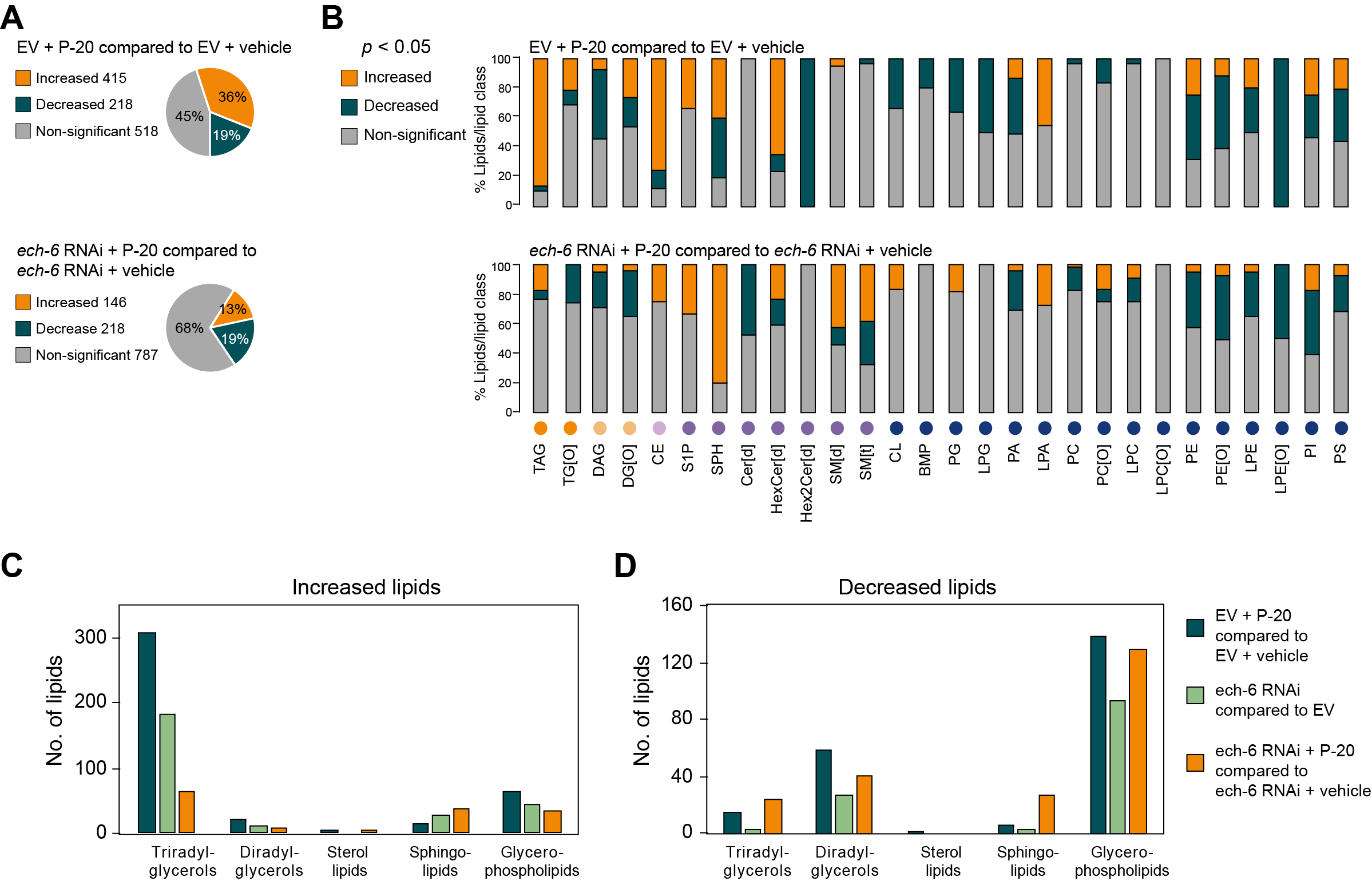

**Figure S2.** Knockdown of *ech-6* Diminishes the Effects of P-20 Supplementation on TAG but Not on Glycerophospholipid Profiles

(A) Pie charts depicting the percentages of increased and decreased lipids by P-20 supplementation. A *p*-value < 0.05 was applied for determining statistical significance.

(B) Percentage distribution of significantly changed lipids by P-20 supplementation across the lipid species in empty vector (EV)- and *ech-6* RNAi-treated worms.

(C-D) The number of increased (C) and decreased (D) lipids in each lipid class upon P-20 supplementation or *ech-6* RNAi. Supplementation of P-20 shows reduced effects on triradylglycerols in *ech-6*-silenced worms compared to its effects in empty vector (EV)-treated wild-type worms, whereas other lipid classes are altered by P-20 supplementation to a similar extent in empty vector (EV)- and *ech-6* RNAi-treated worms. $\chi$^2^ tests, *p =* 0, *p =* 0.1573, *p =* 0.5708, *p =* 0.2846 , and *p =* 0.022 when the number of lipids changed by P-80 supplementation including triradylglycerols, diradylglycerols, sterol lipids, sphingolipids, and glycerophospholipids in empty vector (EV)-treated worms was respectively compared to the corresponding lipid number in the context of *ech-6* deficiency.

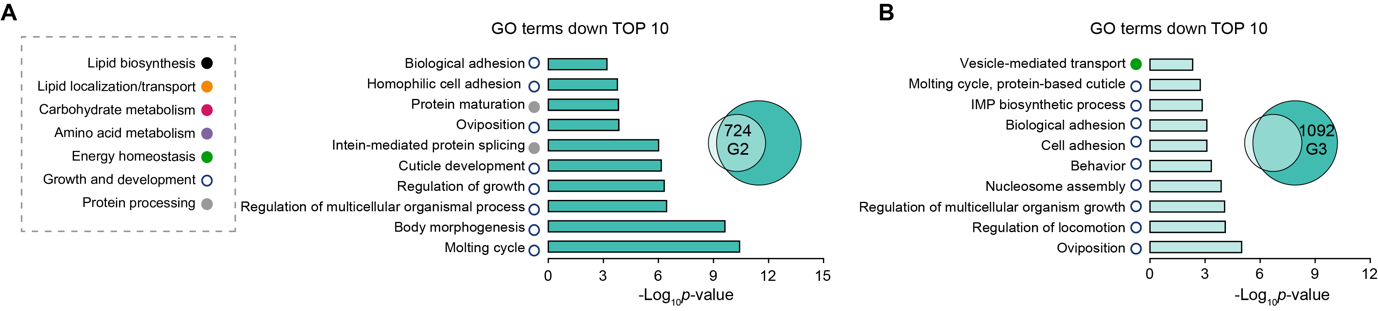

**Figure S3.** Gene Ontology Enrichment Analysis of Downregulated Genes by High-fat Feeding

(A-B) Downregulated genes by P-80 supplementation present in empty vector (EV)- and *ech-6* RNAi-treated worms (G2) and exclusively present in empty vector (EV)-treated worms (G3) are primarily enriched in growth and development-associated processes. GO terms were considered significantly enriched for a modified Fisher’s Exact *p*-value < 0.05 (an EASE score).

**Table S1.** Lifespan Statistics

Summary of median lifespan and statistical analysis (*p*-values) for lifespan experiments displayed in Figures 1B and 1C, Figures 2A and 2B, Figures 2E-2H, Figure 6E. Larval stage 4 (L4) is considered as day 0 of each lifespan assay. The median and *p*-values were calculated by a log-rank (Mantel-Cox) statistical test. *P*-values less than 0.05 are considered statistically significant, demonstrating that the two lifespan populations are different. Statistics of individual experiments are shown for each condition. The number of individuals scored, and independent experiments are shown. a: versus *ech-6* RNAi in N2; b: versus EV + 200 µM P-80; c: versus *ech-6* RNAi + 100 µM P-80 in N2.

**Table S2**. Statistical analysis of lipidomics

**Table S3**. Statistical analysis of RNAseq data
